# Background radiation impacts human longevity and cancer mortality: Reconsidering the linear no-threshold paradigm

**DOI:** 10.1101/832949

**Authors:** Elroei David, Marina Wolfson, Vadim E. Fraifeld

## Abstract

**BACKGROUND:** The current linear-no-threshold paradigm assumes that any exposure to ionizing radiation carries some risk, thus every effort should be made to maintain the exposures as low as possible. Here, we examined whether background radiation impacts human longevity and cancer mortality.

**METHODS:** Our data covered the entire US population of the 3139 US counties, encompassing over 320 million people. The data on background radiation levels, the average of 5-year age-adjusted cancer mortality rates, and life expectancy for both males and females in each county, was extracted using publicly available tools from official sources, and analyzed with JMP^®^™ software.

**RESULTS:** We found for the first time that life expectancy, the most integrative index of population health, was approximately 2.5 years longer in people living in areas with a relatively high vs. low background radiation (≥ 180 mrem/year and ≤ 100 mrem/year, respectively; p < 0.005; 95% confidence interval [CI]). This radiation-induced lifespan extension could to a great extent be associated with the decrease in cancer mortality rate observed for several common cancers (lung, pancreas and colon cancers for both genders, and brain and bladder cancers for males only; p < 0.05; 95% CI).

**CONCLUSIONS:** Exposure to a high background radiation displays clear beneficial health effects in humans. These hormetic effects provide strong evidence for re-considering the linear no-threshold paradigm, at least within the natural range of low-dose radiation.

## Introduction

In the early decades of the 20^th^ century, and mainly during World War II, extensive radiobiological studies were preformed, in order to establish a basic radiation protection policy and philosophy.^1^ The concept of tolerance dose was developed and widely accepted, based on two major postulates: 1) a threshold dose exists, which when exceeded may cause harmful effects; 2) even in case of certain exceeding the threshold, a complete recovery from radiation effects is still possible.^1^ Moreover, an extensive experimental study suggested a beneficial effect of moderate radiation levels on the survival and tumor incidence in mice.^2^ Nonetheless, in 1946, Hermann J. Muller, who received the Nobel Prize in Physiology or Medicine for his work on the mutagenic effect of radiation, stated in his inauguration lecture that “there is no escape from the conclusion that there is no threshold dose, and that the individual mutations result from individual “hits”, producing genetic effects in their immediate neighborhood”.^3^ This strongly led to the embracing of the linear no-threshold (LNT) hypothesis which postulated that there is no “safe” level of radiation, and that even an extremely small dose increases the risk for damage, in direct proportion to the dose.^4^ By the middle of the 1960s, the LNT model for cancer risk assessment was generally considered by the scientific community (and subsequently by the policy makers) as the safest approach to the establishment of radiation protection policy and standards.^5^

Yet, the LNT model remains highly controversial. Among the arguments against the LNT model are that it was not based on concrete scientific data, but rather ideologically motivated and politically influenced.^6-8^ Some even claimed a deliberate counterfeit, up to fabrication of the research record.^6,9^ Most recently, the National Council on Radiation Protection and Measurements (NCRP) published a commentary provided by an interdisciplinary group of radiation experts who critically assessed epidemiologic studies on populations exposed to low dose ionizing radiation, and concluded that the existing data does not challenge the LNT model for radiation protection. As the Scientific Committee proclaims: “It is acknowledged that the possible risks from very low doses of low linear-energy-transfer radiation are small and uncertain and that it may never be possible to prove or disprove the validity of the linear no-threshold assumption by epidemiologic means… Currently, no alternative dose-response relationship appears more pragmatic or prudent for radiation protection purposes than the linear no-threshold model.”^10^

In this study, we undertook a comprehensive analysis of the dose-response to background radiation in the entire US population, with regard to life expectancy and site-specific cancer mortality.

## Methods

### DATA SOURCES

Our data included the entire US population of the 3139 US counties, encompassing a total number of over 320 million people. The variables on the analysis included the background radiation levels, age-adjusted cancer mortality rates, and life expectancy data for each county. Data on radiation levels was calculated using the United States Environmental Protection Agency’s (EPA) radiation dose calculator (https://www.epa.gov/radiation/calculate-your-radiation-dose), taking into account the two major sources of background radiation: terrestrial radiation and cosmic radiation. Information about specific average height of every county was taken from The National Map (TNM) tool of the United States Geological Survey (USGS) (https://viewer.nationalmap.gov/theme/elevation/#). Data on cancer statistics was collected from the United States Cancer Statistics (USCS), the official federal statistics on cancer incidence and deaths, produced by the Centers for Disease Control and Prevention (CDC) and the National Cancer Institute (NCI) (https://www.statecancerprofiles.cancer.gov/deathrates/index.php). The collected data included the average of 5-year (2011-2015) age-adjusted cancer mortality rates (total and site-specific) for the entire US counties, separately for males and females. Data on life expectancy was collected from the Institute for Health Metrics and Evaluation (IHME), an independent population health research center at the University of Washington Medical Center (http://ghdx.healthdata.org/us-data).

### DATA ANALYSIS

Data was analyzed using JMP^®^™ software for statistical analysis. The data was screened for outliers, entry errors, missing values and other inconsistencies that could compromise the analysis. Jackknife resampling technique was used to estimate the bias and the variance of the data. Since cancer and life expectancy data exhibited normal distribution (Suppl. Fig. 1), the statistical significance of the differences between the categories was determined by *t*-test or by ANOVA test, when two categories or more were involved, respectively. Data on esophagus, melanoma of the skin, oral cavity & pharynx, thyroid, and uterus cancers was not sufficient enough for a reliable statistical evaluation, and thus was not included in the analysis. Both parametric (Pearson) and non-parametric (Spearman) coefficients of correlation were estimated. In all cases the parametric and non-parametric estimates were in good agreement, thus the results of only parametric analyses were presented. The *p*-values of less than 0.05 were considered statistically significant.

## Results

### DISTRIBUTION OF BACKGROUND RADIATION LEVELS IN THE US

The background radiation levels have a 2.5-fold difference between the lowest and the highest levels, ranging from 92 to 227 mrem/year, with a median value of 115 mrem/year. The lower levels are mainly found at the Gulf and Atlantic coasts states, and the higher levels are mainly found at the Colorado Plateau states (Fig. 1).

**Figure 1:**
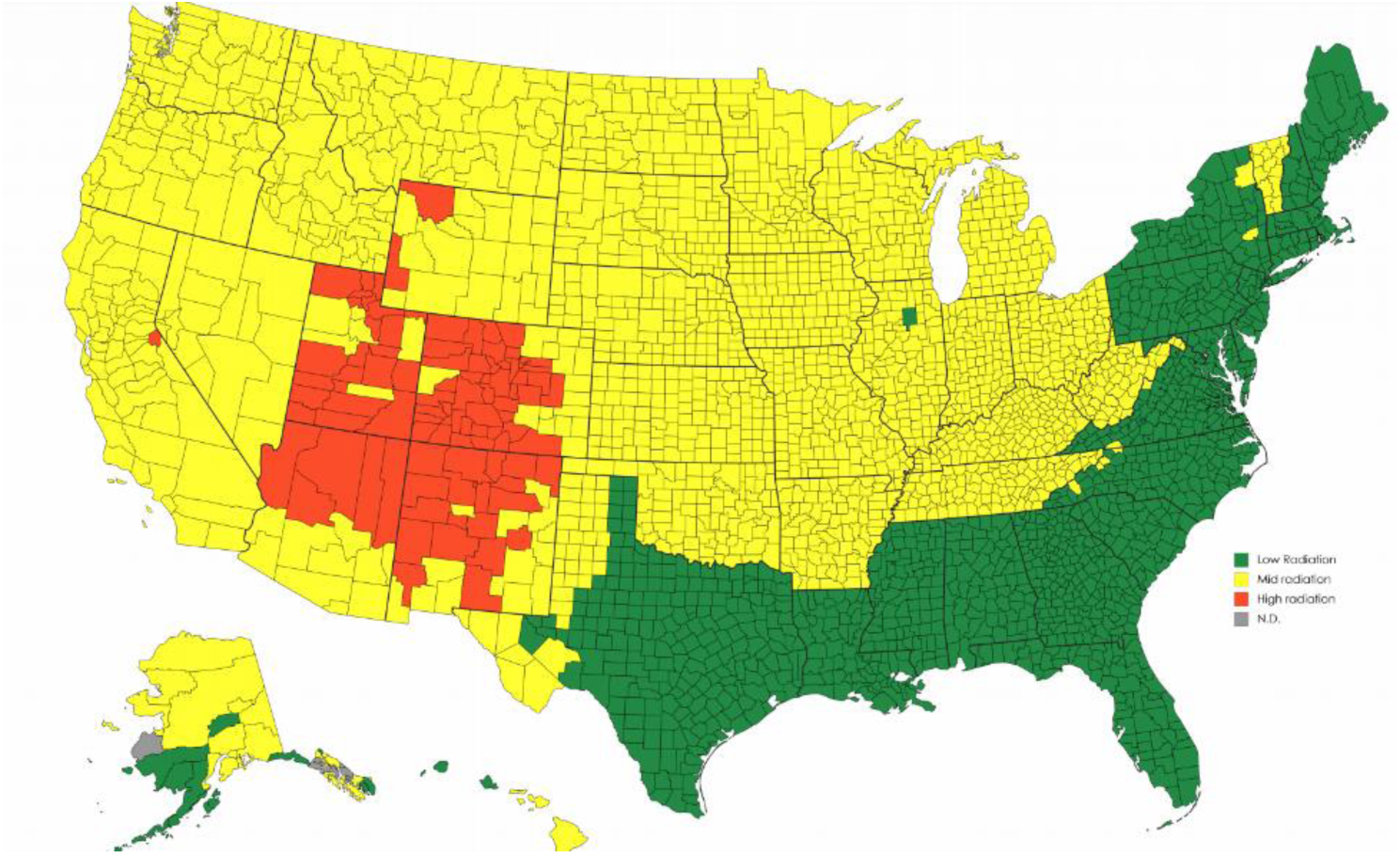
A map of background radiation levels in the US counties. The background radiation levels were divided into 3 categories: the levels lower than 100 mrem/year were defined as “Low radiation”; the levels higher than 180 mrem/year were defined as “High radiation”, and the levels between 100 mrem/year and 180 mrem/year were defined as “Mid radiation”.

### LIFE EXPECTANCY

As seen in Fig. 2, a highly significant positive correlation of life expectancy with background radiation levels was found for both males and females (r = 0.647 and r = 0.651, respectively; p < 0.001; 95% confidence interval [CI]). According to the regression line, every 10 mrem/year extend the life expectancy by 2.4 months in males, and by 1.8 months in females, so that within the natural range of background radiation this provides a maximum increase in life expectancy of 2.7 and 2.4 years for males and females, respectively.

**Figure 2:**
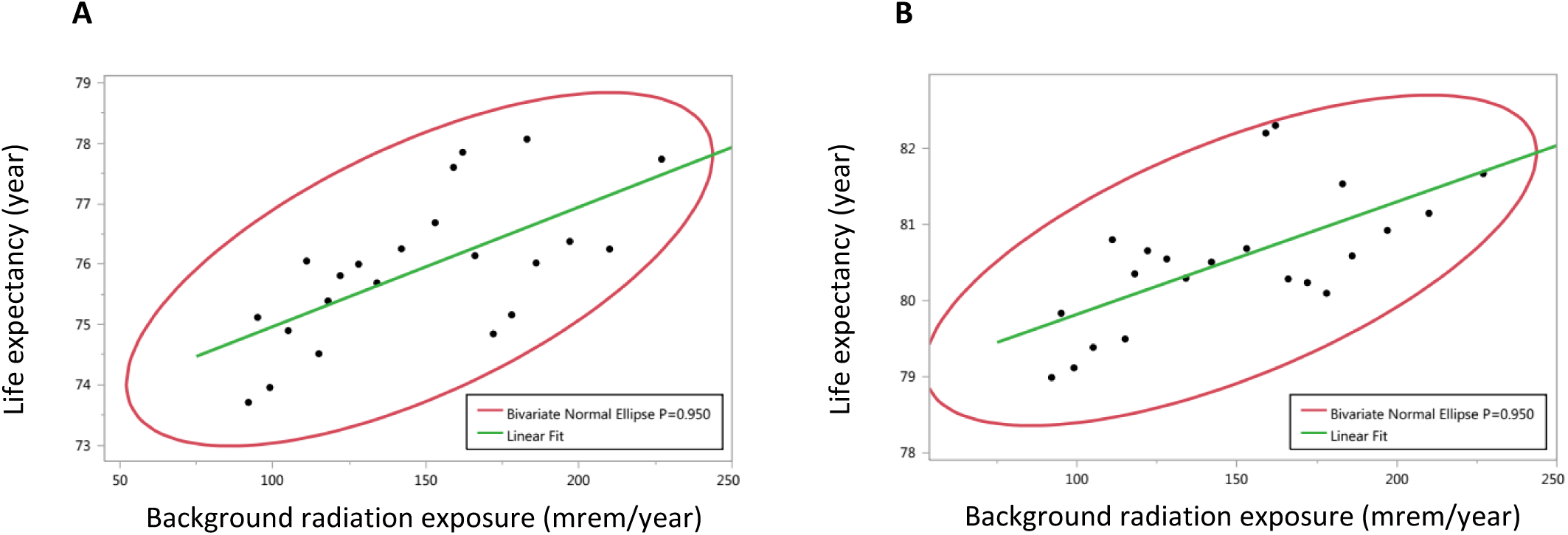
The life expectancy was plotted against the average estimated radiation level in each county. The regression equations for males (**A**) and females (**B**), respectively: *LE* = 0.02*BRL* + 73.0 and *LE* = 0.015*BRL* + 78.3, where LE stands for life expectancy in years, and BRL stands for background radiation levels in mrem/year.

### TOTAL CANCER MORTALITY RATE

The analysis revealed a strong negative correlation between age-adjusted cancer mortality rates and background radiation levels, for both males and females (r = -0.90 and r = -0.77, respectively; p < 0.001; 95% CI). According to the regression line, an increase in background radiation by approximately 2 mrem/year in males and 4 mrem/year in females decreases the total cancer mortality rate by 1 death per 100,000 people. Thus, within the natural range of background radiation, the maximum radiation-dependent effect would be around 69 deaths per 100,000 people in males (p < 0.001; 95% CI), but almost two-fold lower in females (35 deaths per 100,000 people; p < 0.001; 95% CI) (Fig. 3).

**Figure 3:**
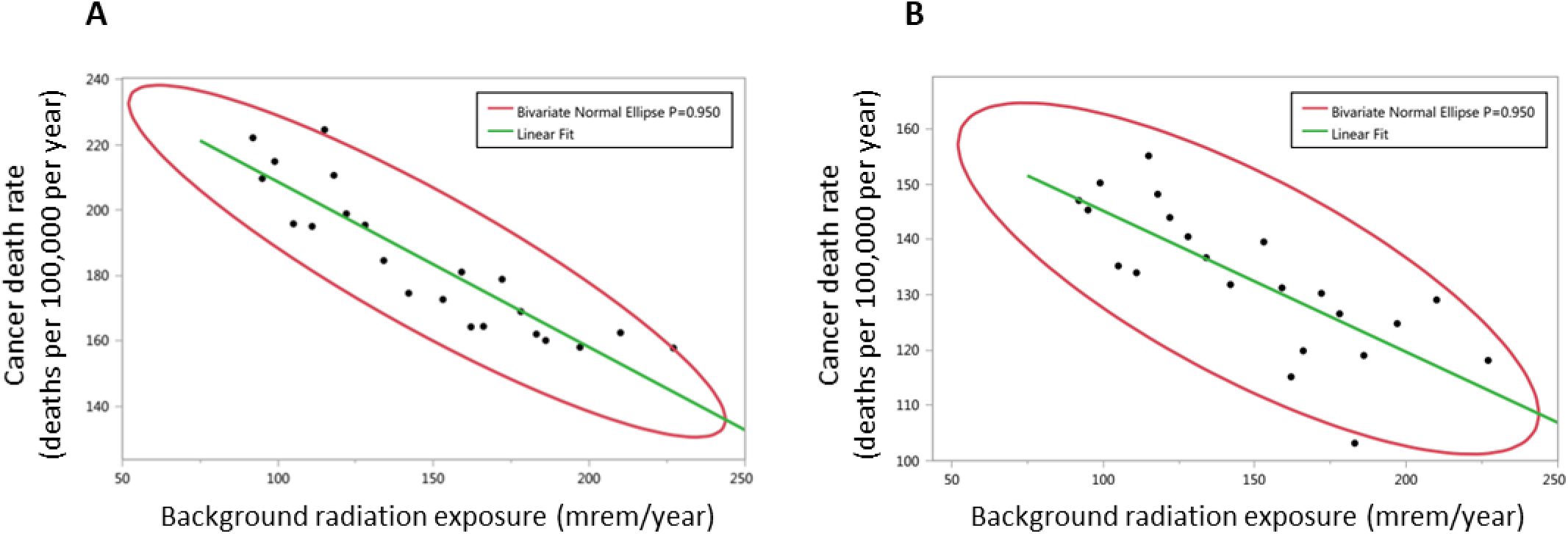
Background radiation levels were plotted against age-adjusted cancer mortality rates. The regression equations for males (**A**) and females (**B**), respectively: *CDR* = −0.51*BRL* + 259.2 and *CDR* = −0.26*BRL* + 170.7, where CDR stands for the number of cancer deaths per 100,000 per year, and BRL stands for background radiation levels in mrem/year.

### RADIATION EFFECT ON SITE-SPECIFIC CANCER MORTALITY RATES

Our next step was to evaluate whether the radiation-dependent effects on age-adjusted cancer mortality rates are site-specific. For this purpose, we compared the effects of low vs. high background radiation levels, as categorized above (see Fig. 1). The results of this analysis are presented in Table 1. In both males and females, we discovered a significant decrease (p < 0.005; 95% CI) in mortality rates for lung & bronchus cancer, pancreas cancer, and colon & rectum cancer. In males, but not in females, a significant decrease for brain & ONS cancer, and bladder cancer (p < 0.05; 95% CI) was also observed, as well as a clear tendency toward significance for liver & bile duct cancer (p = 0.08; 95% CI). In contrast, neither for males nor for females, any significant effects were found for leukemia, kidney & renal pelvis cancer, and stomach cancer (p > 0.2; 95% CI), as well as for the gender-dependent cancers (cervix, breast, and prostate; with a clear tendency for ovarian, p = 0.08; 95% CI).

**Table 1.** Site-specific cancer mortality rates (deaths per 100,000 people; the means ± SD) in response to low-level radiation (LLR) and high-level radiation (HLR), in both males and females.

| Cancer type | Males |  | Females |  |
| --- | --- | --- | --- | --- |
|  | LLR | HLR | LLR | HLR |
| Lung & Bronchus | 68.6 ± 18.8 | 42.5 ± 20.3 * | 41.1 ± 10.7 | 30.7 ± 11.7 * |
| Pancreas | 13.8 ± 3.2 | 12.2 ± 3.2 * | 10.2 ± 2.4 | 9.5 ± 1.8 * |
| Colon & Rectum | 20.4 ± 6.1 | 18.6 ± 6.0 * | 13.8 ± 3.7 | 12.1 ± 3.7 * |
| Brain & ONS | 5.8 ± 1.6 | 5.1 ± 0.7 * | 3.9 ± 1.1 | 3.7 ± 0.5 |
| Bladder | 8.4 ± 2.1 | 7.5 ± 1.7 * | 2.4 ± 0.6 | 2.3 ± 0.3 |
| Liver & Bile duct | 10.2 ± 3.0 | 9.1 ± 3.6 ** | 4.0 ± 1.4 | 4.3 ± 1.3 |
| Breast | - | - | 22.6 ± 5.1 | 21.9 ± 6.1 |
| Leukemia | 9.5 ± 2.1 | 9.6 ± 2.8 | 5.3 ± 1.2 | 5.1 ± 1.1 |
| Kidney & Renal pelvis | 6.0 ± 1.6 | 6.1 ± 1.8 | 2.6 ± 0.8 | 2.5 ± 0.6 |
| Cervix | - | - | 2.5 ± 0.8 | 2.4 ± 0.7 |
| Ovary | - | - | 7.6 ± 1.8 | 8.1 ± 1.3 ** |
| Prostate | 21.8 ± 6.6 | 22.2 ± 3.9 | - | - |
| Stomach | 4.8 ± 1.5 | 5.4 ± 2.7 | 2.5 ± 0.7 | 2.5 ± 0.8 |

\* $p < 0.05$ ; \*\* $p = 0.08$

## Discussion

The linear no-threshold (LNT) model of radiation damage is still widely accepted.^10,11^ Moreover, this concept determines the current radiation protection policy as reflected by the ALARA guiding principle of the International Commission on Radiological Protection (ICRP): “<u>A</u>s <u>L</u>ow <u>A</u>s <u>R</u>easonably <u>A</u>chievable”.^12,13^ ALARA assumes that any exposure to ionizing radiation carries some risk, thus every effort should be made to maintain the exposures *as low as possible*. Consequently, each year hundreds of billions of dollars are spent worldwide to maintain extremely low radiation levels.^14^ Yet, our results provide strong evidence for *re-considering* the LNT paradigm, at least within the natural range of background radiation. Indeed, we have shown that not only that the highest background radiation levels (from 180 and up to 227 mrem/year) do no harm compared to the lowest levels of less than 100 mrem/year, but also display clear beneficial health effects. Life expectancy, the most integrative index of population health, was found to be approximately 2.5 years longer in people living in areas with a relatively high vs. low background radiation. This radiation-induced lifespan extension could to a great extent be associated with the decrease in cancer mortality rate observed in HLR areas for several common cancers including lung, pancreas, colon, brain, and bladder cancers.

As mentioned above (see Introduction), the NCRP experts recently re-evaluated the LNT model for radiation protection policy and came to the conclusion that there is no reason to modify the current concept.^10,15^ This conclusion may appear controversial to our findings on the longevity-promoting and cancer mortality-reducing effects of high-level background radiation. However, a more careful examination of the data reveals otherwise. First, the range of radiation levels analyzed in the NCRP’s study was at least several folds higher (and in many cases even several orders higher) than in our study. Second, even in those relatively high levels, no significant association was found between radiation and several site-specific cancers, among which are gender-dependent cancers (cervix, breast, and prostate) and leukemia^10^ — the results that are in fact consistent with ours. Furthermore, the NCRP’s study did not relate to cancers in which we did find a significant negative correlation of mortality rates with background radiation levels (i.e., pancreas, colon, brain, and bladder). All in all, it is reasonable to suggest that a radiation threshold does exist, yet it is higher than the upper limit of the natural background radiation levels in the US (227 mrem/year). Below the threshold level, the opposed relationships between the background radiation and its health effects are observed, so that the higher radiation exposure, the longer life expectancy and the lower cancer-associated mortality.

While the biological effects of high radiation (generally artificial) were extensively investigated,^13,16,17^ the studies on the effects of low levels (comparable with natural background) on human health and longevity are limited, and often inconclusive because of the relatively small population size analyzed. Overall, these studies, which are reviewed elsewhere,^18-20^ are in line with our major findings on the association of higher background radiation with less cancer mortality and lifespan extension. Experimental studies on mammals (mice, rats, beagle dogs, chipmunks, etc.) also suggest the existence of a radiation range with lifespan-extending effect, a phenomenon known as “longevity hormesis”.^2,21-23^ Hormetic effects of low-dose radiation could be the result of several potential mechanisms, such as the activation of DNA repair, activation of endogenous antioxidant systems, induction of the heat shock protein response, stimulation of immune responses, etc.^24-27^ Whatever the case, it seems that Kondo was indeed correct when stated that “*The collected data strongly suggest that low-level radiation is not harmful, and is, in fact, frequently ‘apparently beneficial’ for human health.”*^28^

